# Two-Dimensional Mapping of 3D Data from Confocal Microscopy

**DOI:** 10.1101/019133

**Authors:** Anthony Fan, Justin Cassidy, Richard W. Carthew, Sascha Hilgenfeldt

**Affiliations:** Department of Mechanical Science and Engineering, University of Illinois, Urbana, IL, 61801 USA; Department of Molecular Biosciences, Northwestern University, Evanston, IL, 60208 USA

**Keywords:** Confocal Microscopy, Image Reconstruction, Geodesics, 2D Mapping

## Abstract

Confocal microscopy has been experimentally proven for decades to provide high-quality images for biological research. Its unique property of blocking out-of-focus light enables 3D rendering from planar stacks and visualization of internal features. However, visualizing 3D data on a flat display is not intuitive, and would lead to occasional distortion. In this study, a novel, easy-to-implement, and computationally fast solution is provided to reconstruct a confocal stack to true 3D data and subsequently map the information correctly with size and shape consistency to a 2D space for visualization and image analysis purposes.

## I. INTRODUCTION

Confocal microscopy provides high resolution images with almost no out-of-focus blur [1]. The specimen is usually tagged with fluorescing molecules or is itself autofluorescing. Together with a scanning laser source of small Spot size, it can easily provide sub-micron spatial resolution. Because of the advantage to see only in focus features, confocal imaging is particularly useful in obtaining 3D information and has been widely used for such.

Bio-imaging in the recent decades has attracted a lot of attentions, and confocal imaging has proven to be the prime tool for such applications [2], [3]. It has been shown successful in *in vitro*, *in situ*, and *in vivo* imaging ranging from the sub-cellular level [4] to the tissue and small animal level [5]. Confocal is particularly beneficial in bioimaging because minimal treatment is required on delicate biomaterials which also implies the possibility for live imaging. Blocking of out-of-foucs light also enables visualizations of internal features within the focal plane that will be blurred by scattering contamination in traditional light microscopy. Combining with a motorized lens or stage, in which the z-axis actuation is accurately controlled, the confocal microscope can provide z-stack images spanning across the entire specimen, if the working distance allows, allowing 3D visualization.

3D reconstruction is mature and most confocal acquisition softwares have such tools built-in. In the simplest form, it is the projection of maximum intensity along the viewing axis. As the viewing angle is altered, the projection is done in different direction thus enabling 3D rendering to the user. However, it is not true 3D information, and limit interpretation to largely qualitative. Many efforts have been demonstrated in presenting the 3D information in a variety of systems [6], [7], [8], [9]. Here, I will use the compound eye of *drosophila* as an example to present a method to accurately reconstruct a confocal stacks into a point cloud form, with each point tagged with an intensity (4D) value. However, visualization of such data on any form of display (which is 2D) is still non-intuitive, therefore I present a method based on geodesics estimation to reconstruct such 3D data back to a 2D plane without size and morphology distortion from uniaxial projection.

## II. ANIMAL MODEL, IMAGING, AND ENABLING SOFTWARE

Room temperature red-eye wild-type *drosophila* were used. All flies imaged were sacrificed using an ethanol step-dry process to preserve eye geometry and to enable SEM imaging for confirmation. Confocal imagings were done in the Materials Research Laboratory (MRL) at the University of Illinois using the Zeiss LSM 7 Live. SEM imagings were done using Hitachi 6060LV also in MRL. All image post-processing algorithms were written in MATLAB. Some minor visualizations and boundary trimming were done in ImageJ.

## III. CONFOCAL STACKS TO 3D POINT CLOUD

A confocal stack can be described by the function *I*[*x, y, z*], in which every spatial coordinate will have an intensity value. Each *x*-, *y*–coordinate gives rise to a unique *I*_*x,y*_[*z*] function (Fig. 1A). This is expected to be approximately Gaussian because of remaining out-of-focus blur. Therefore, a two-term Gaussian fit is performed on all *I*_*x,y*_[*z*] from which the fitted maximum intensity, *I*_*max*_[*x, y*], and the corresponding z-position, *zI*_*max*_[*x, y*], could be obtained. This leads to a 3D point cloud surface that could be subsequently described mathematically (Fig. 1B).

**Fig. 1.**
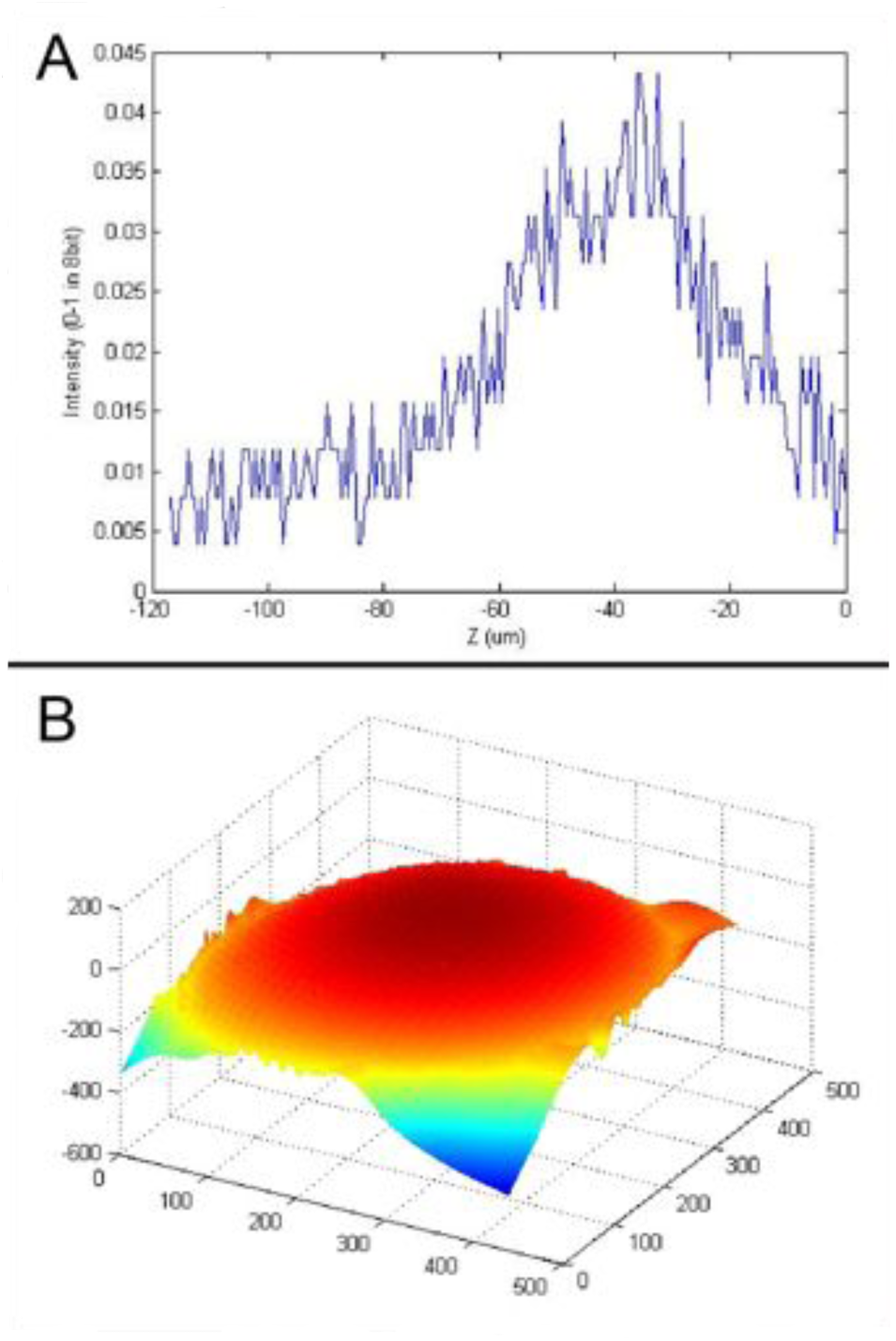
(A) Intensity vs. *z*-position profile at a randomly chosen *x*-, *y*-coordinate. (B) The constructed 3D point cloud matched to a surface.

## IV. FITTING OF 3D POINT CLOUD

The 3D data is then fitted to an ellipsoid:

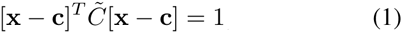

or in polynomial form:

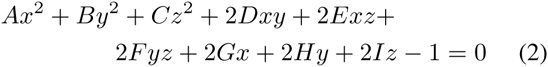

one can characterize the surface exactly with 9 degrees of freedom: 3 translations, 3 rotations, and 3 semi-principal axes. It is important to note that other, or a combination of, mathematical structures might be more suitable depending on the application. Here, for demonstration purposes, I find a general ellipsoid adequate.

By evaluating the error spatially, one can confirm that the ellipsoid is in close agreement with the 3D data. Therefore, the distance between each point with the surface of the ellipsoid is calculated and plotted over the best-fit ellipsoid in Fig. 2A. The error distribution is shown in Fig. 2B. As apparent, the fitting error is small comparing to the magnitude of the semi-principal axes (subplot of Fig. 2B), which gives confidence in proceeding to the next section in performing 2D mapping.

**Fig. 2.**
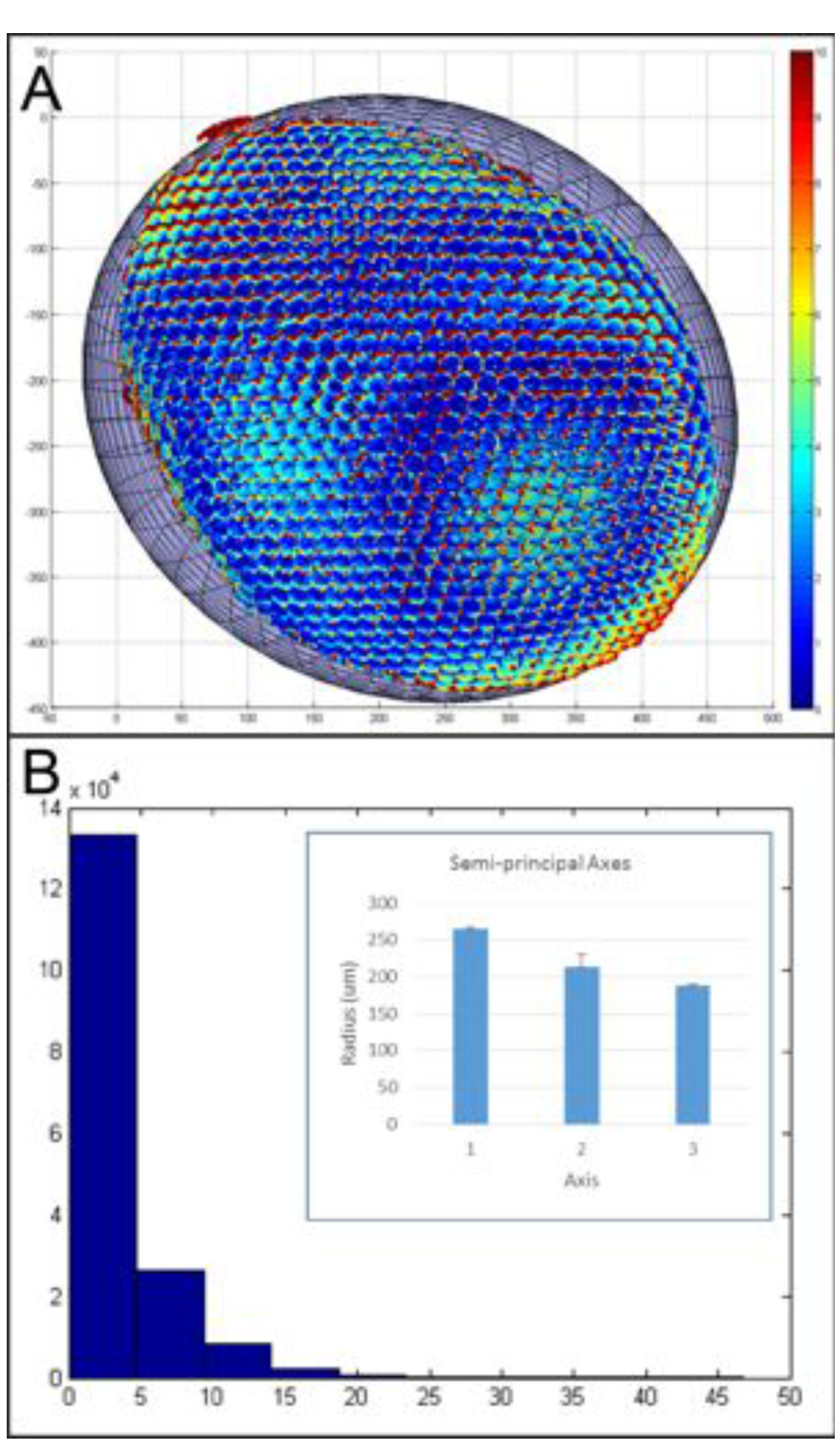
(A) Raw data points on best-fit ellipsoid. Error magnitudes are shown in color with blue as close to *±*±0 *µ*m and red as *±*±10*µ*m or above. (B) Error distribution with *x*-axis in unit of *µ*m and *y*-axis as bin frequency. Most points have error less than 5 *µ*m. Subplot shows the magnitude of semi-principal axes in best-fit ellipsoid. Error is small compared to the overall geometry. Error bars in SD. N=2.

## V. MAPPING 3D POINT CLOUD TO 2D IMAGE USING AN ELLIPSOID FIT

### A. Center as Reference and Point Closest to the Camera as Point of Projection

The undistorted point of projection is the maximum point along the **e_3_** direction as depicted in Fig. 3. Differential geometry enables explicitly obtaining the coordinate of maximum z(*maxz*) by locating the zero tangent plane:

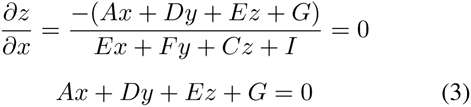

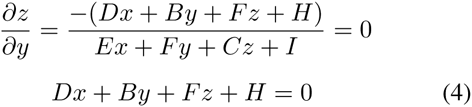

solving equation (2), (3), and (4) will lead to a close form solution for *maxz* = [*x*_*maxz*_, *y*_*maxz*_, *z*_*maxz*_].

**Fig. 3.**
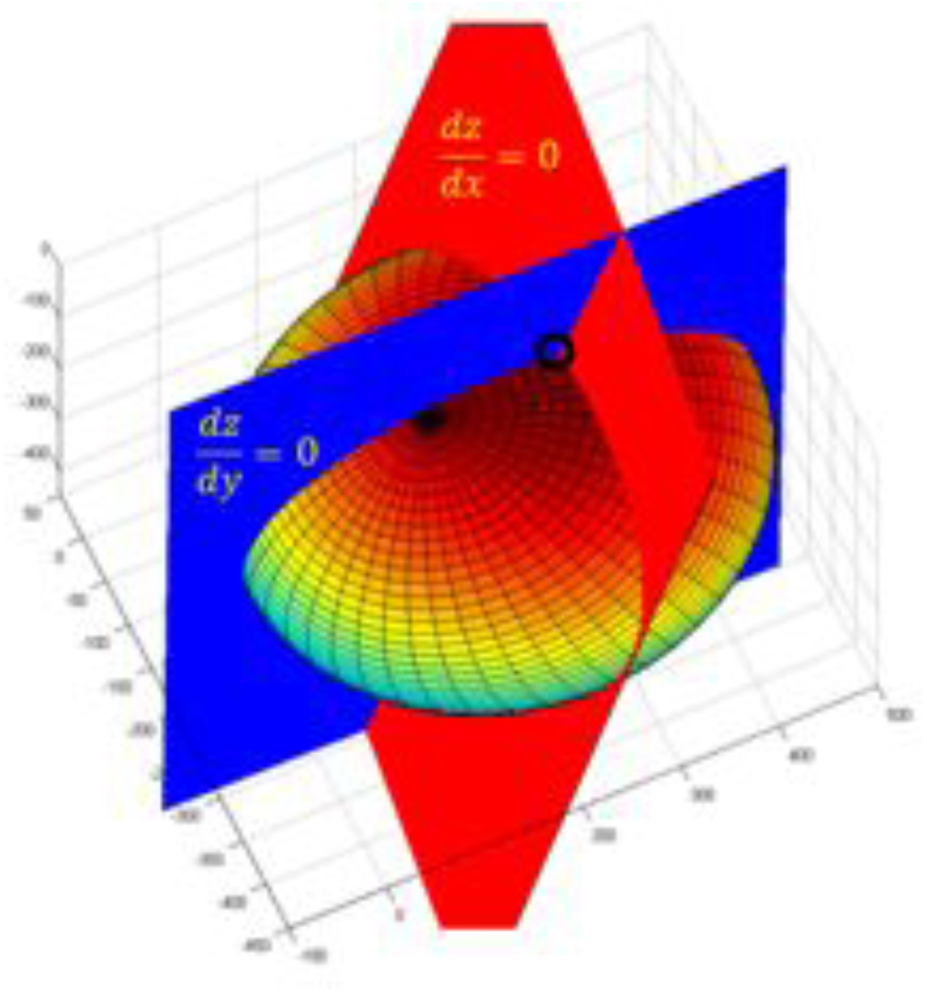
Visualization of the point *maxz* (black circle): intersection of the plane 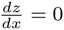 (red), the plane 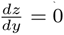 (blue), and the ellipsoid.

A dissection plane (Fig. 4) can then be constructed from 1) the vector, **s**, connecting *maxz* and the center of the ellipsoid (*center*) and 2) the vector, **t**, connecting *maxz* and any point on the surface of the ellipsoid (*point*). This plane cuts through the center, the projection reference i.e. *maxz*, and the point of interest.

**Fig. 4.**
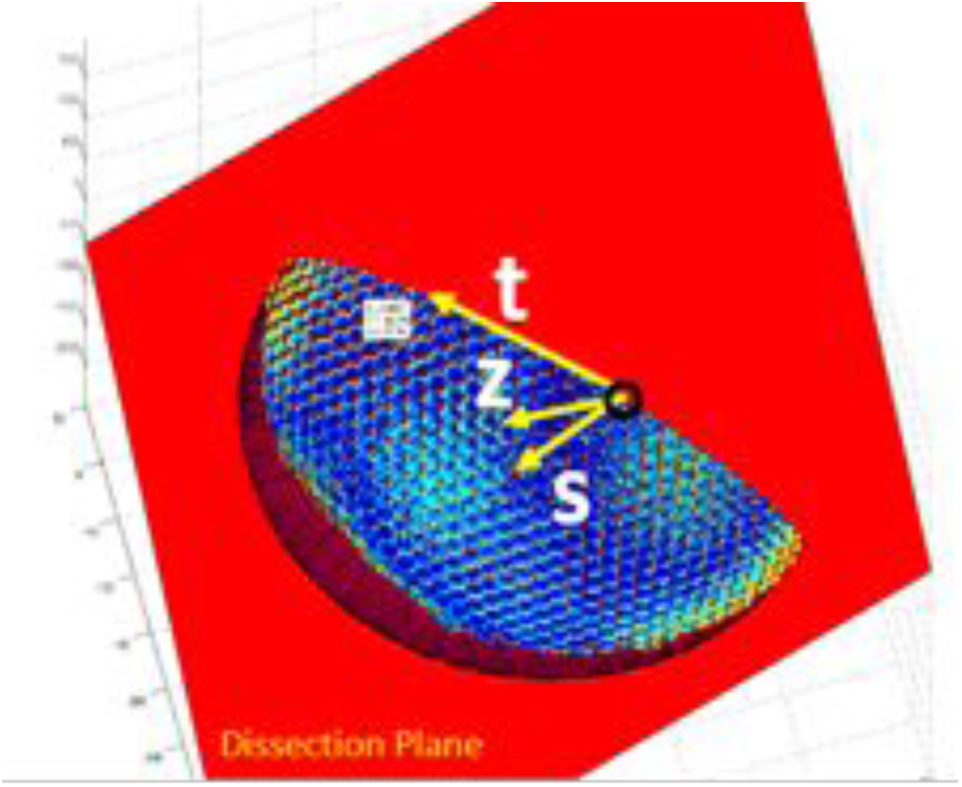
Construction of the dissection plane for a randomly selected point. **z** is the plane normal. **t** is the vector connecting *maxz* and *point*. **s** points to *center* from *maxz*.

The plane normal, **z**, is defined as:

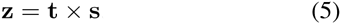

Using the plane normal in Eq. 5, one can construct the complete rotational matrix describing the orientation of the plane by obtaining the vector, **y**, mutually orthogonal to **s** and **z**, where

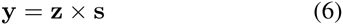

And thus, the rotational matrix is in the form of

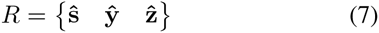

where 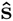, 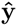, and 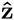 unit column vectors of **s**, **y**, and **z**.

*R* is used to align the dissection plane with the *z* = 0 plane. Such alignment enables path integration from *maxz* to *point* to compute the arc length between the 2 points, providing a quick and accurate estimate of the geodesics.

The alignment process is as follows. The center of the ellipsoid could be translated to the origin and its formula is then:

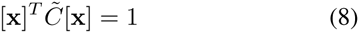

If no rotation is imposed, 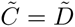 in Eq. 8 is a diagonal matrix containing the semi-principal axes information. If a rotation *R*_0_ is imposed, 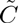 in Eq. 8 is a symmetric matrix in which:

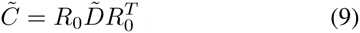

Now, with the previously calculated *R* from Eq. 7, we can formulate a new 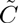, 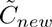:

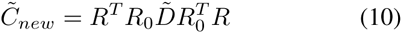

and the form of the ellipsoid is then

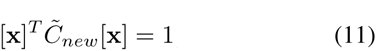

In which substituing *z* = 0 into Eq.11 will give the ellipse connecting *maxz* and *point* with center = origin as the center of the ellipse. Note that *maxz*, *point*, and *center* will not have the same coordinates as before because of translation and rotation. Their distances and relative angles, however, are conserved, given by the properties of rotational matrices.

The estimated geodesics (Fig. 5), *Geo*, can then be obtained by doing a path integral from *maxz* to point:

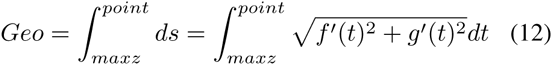

**Fig. 5.**
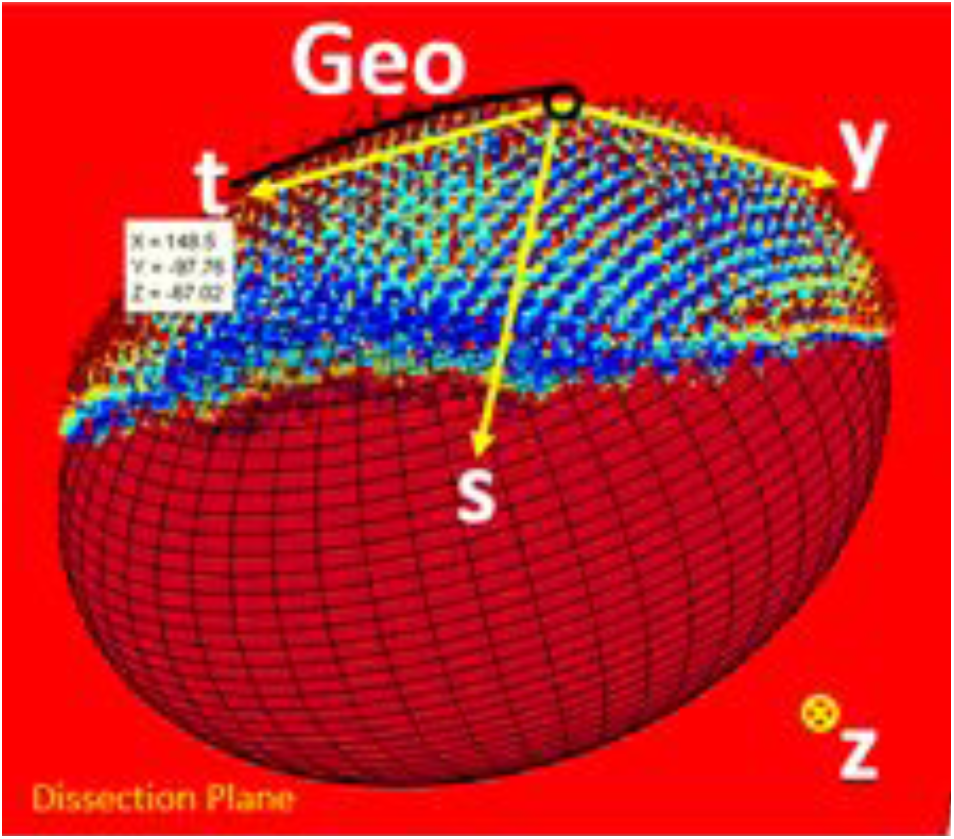
The estimated geodesics is the arc length of the arc connecting *point* and *maxz* along the surface of the ellipsoid now aligned to the plane *z* = 0.

Where *f*(*t*) is the x-parametrization and *g*(*t*) is the y-parametrization of the ellipse. Parametrization guarantees the uniqueness of solution. F(t) and g(t) takes the form:

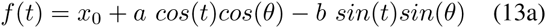

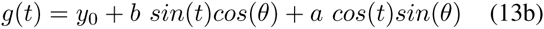

where *θ* is the angle of rotation of the semi-principle axes *a* and *b*.

The intersection of the dissection plane and the plane *z* = *z*_*maxz*_ will give the direction of reconstruction. *Geo* will give the distance (Fig. 6). Explicitly,

**Fig. 6.**
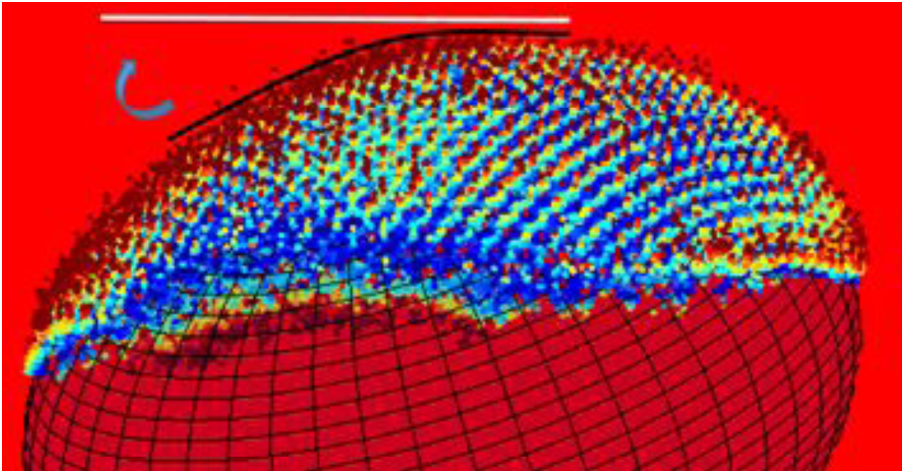
The estimated geodesics is used as the projection distance along the line describe by the intersection of *z* = *zmaxz* and the dissection plane. Every point will get a new reconstructed coordinate this way.

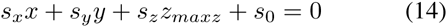

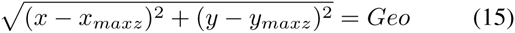

where *s*_0_ is the plane constant. Solving Eq. (14) and (15) will yield the reconstructed *x*-, *y*-coordinates.

Reformatting the x-, y-coordinates and respective maximum intensity value, *I*(*x, y*), back to an image is achieved by weighted interpolation of each pixel (*x*_*i*_,*y*_*i*_) with the nearest N points:

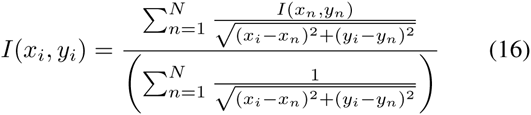

This search of the nearest N-points in Eq. 16 is based on a previously developed algorithm [10]. The author finds that a 4-point interpolation yields the best result.

### B. Plane Normal as Reference and any Point on Surface as Point of Projection

In the previous subsection, we used the cendter of the ellipsoid as a reference for the dissection plane. But some structure do not have a natural center and it is easier to use the normal of the plane tangent to the point of projection (maxz in the previous case) as the reference instead.

In some occasions, the user might want to perform the projection at a point other than the one closest to the camera. The tangent plane normal in these instances will have components other than **e_3_**. The projection therefore will have to be done by also considering the z-dimension. This plane can be subsequently rotated back to align with the x-y plane for image formulation as desribed.

Using the plane normal as reference, the dissection plane does not cut through the center (unless the plane normal passes through the center, which is rare), and, therefore, the arc length is not part of an ellipse that can be simply described by 2 semi-principle axes with a rotation as in Eq. 13. In our reconstruction we will assume that it is such an ellipse regardless, by matching the magnitude and rotation of the semi-principal axes to the curve:

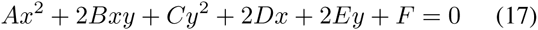

Eq. 17 is the most general form of an ellipse when *B*^2^ *< AC*, as in what would appear when a plane intersects a triaxial ellipsoid arbitrarily. But the system is over-determined with 6 variables–the most general regular ellipse would have 5 variables: 2 semi-principal axes, 2 variables for the center location, and 1 in-plane rotation. The authors are not aware of the parametric form of Eq. 17, and for computation speed and uniqueness considerations proceed with the approximate parametrization, by assuming a regular ellipse.

## VI. MAPPING 3D POINT CLOUD TO 2D IMAGE WITHOUT OBJECT FITTING

To extend the generality of the method. We explore reconstruction without assuming the eye as a triaxial ellipsoid.

First a convex hull is constructed to the 3D point cloud data after section III. This is achieved first by smoothing the data and then by employing the Qhull algorithm [ref] to triangulate the surface.

Once the convex hull is constructed. The geodesics distances and initial angle of the geodesic paths from the reference point to all the vertices are numerically calculated (Fig. 7) [ref]. With the distances and the initial angles, we can map the 3D vertices from (*x*_0_, *y*_0_, *z*_0_) onto the same plane *P* as (*x*_*p*_, *y*_*p*_).

**Fig. 7.**
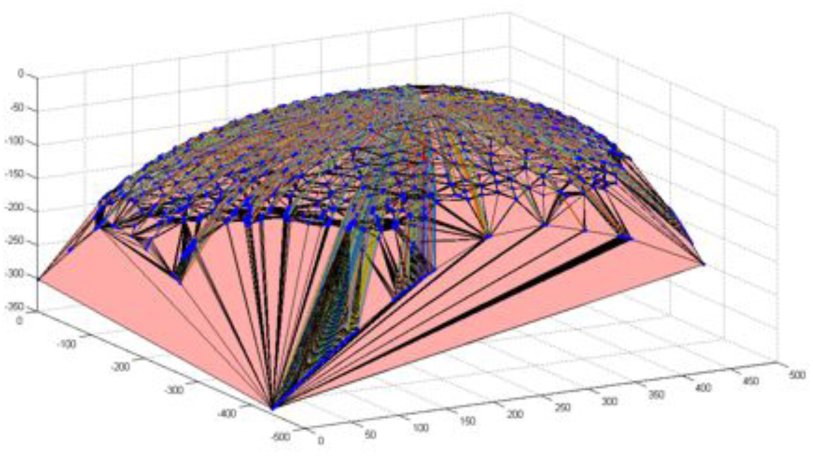
Geodesics path from point of projection to all vertices of convex hull.

The triangulation creates piecewise planar surfaces that tile the curved surface. Intensity projection creates projection points (*x*_*q*_, *y*_*q*_) on the x-y plane *Q*. Each planar surface uniquely contains a set of projection points from *Q*, which could be described by 2 bases. These 2 bases are formulated by the 2 vectors, *v*_1*q*_ and *v*_2*q*_, connecting any one vertice of the triangle to the other 2 vertices on *Q*. This is chosen specifically this way because we know the vertices, as well as the vectors, are mapped *Q → P* and the remaining points in *Q* could be linearly transformed onto *P* by:

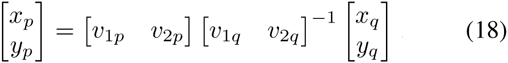

To determine if a projection point from Q is contained in the planar triangle, 3 criteria has to be simultaneously satisfied: 1) *m >* 0, 2) *n >* 0, and 3) *m* + *n <* 1, where:

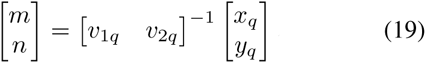

The rest of the process is the same as in section V(A).

## VII. RECONSTRUCTED IMAGES

The original maximum intensity projection image is shown in Fig. 8A and the reconstructed image using the ellipsoid center as reference is shown in Fig. 8B. These pictures come from the compound eye of a *drosophila*. Each individual eye is called an ommatidium. As can be identified, the correction to size and morphology is significant. Especially at the lower right boundary, where previous ommatidia are distorted and cannot be nicely identified. After correction, all ommatidia is of consistent shape and size throughout the entire image, expect where the fit deviates at the left boundary (Fig. 2A). SEM images (Fig. 8C) are taken to show that boundary ommatidia should have size and morphology consistent with bulk ommatidia. It is even more obvious upon closer investigations revealing changes before and after reconstruction (Fig. 8D).

**Fig. 8.**
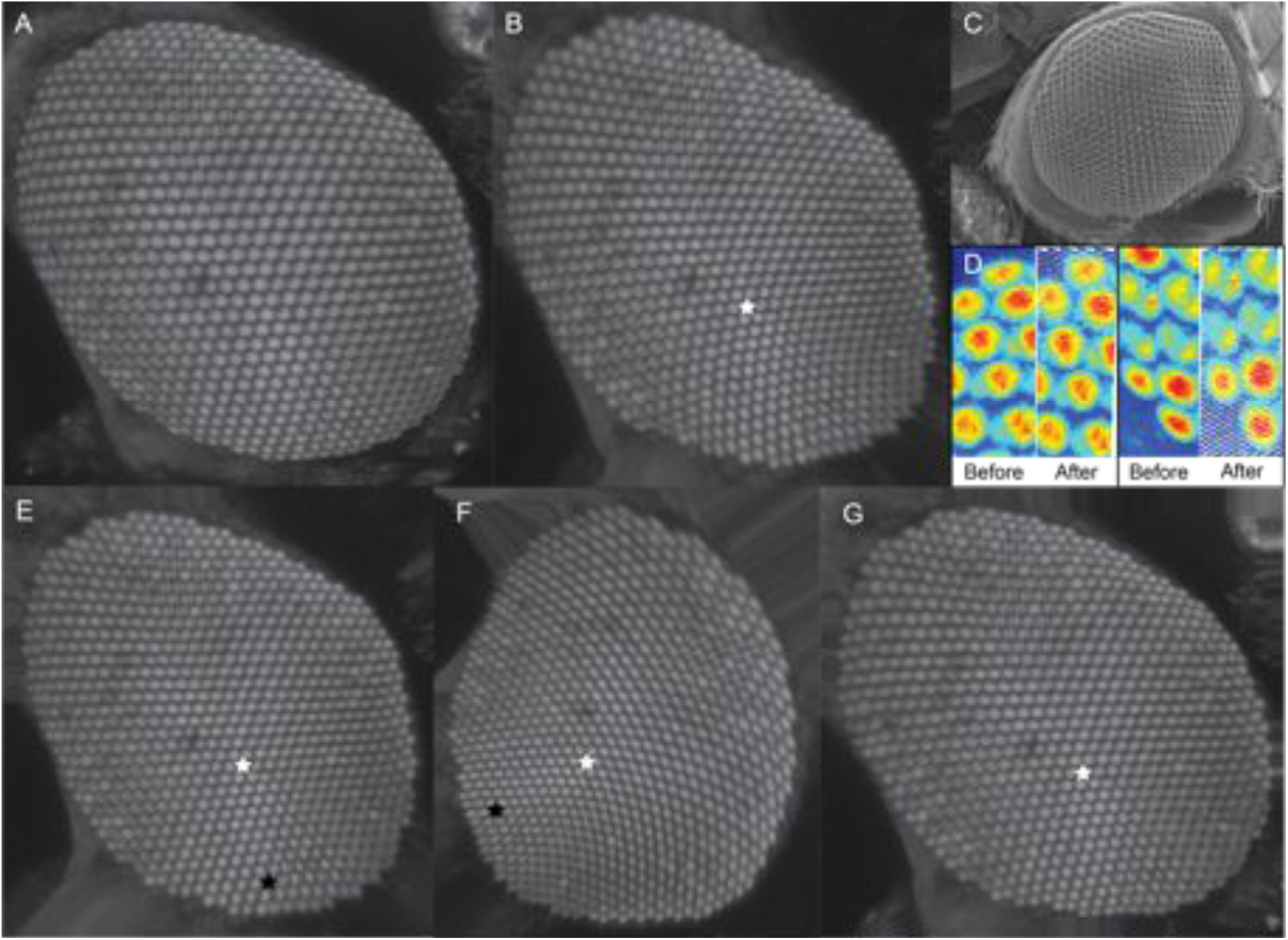
(A) Original image from a maximum intensity projection. (B) Reconstructed image using the image in Fig. 8A with the algorithm described in section V(A). Notice that the ommatidia at the boundary now have corrected size and morphology. Distortion from uniaxial projection is minimized. (C) SEM image taken from a tilted angle to confirm that boundary ommatidia have similar size and shape compared to bulk ommatidia. (D) Close-up view of 2 occasions of boundary ommatidia. Images shown here are before 4-point interpolation to demonstrate the survey of the reconstruction. (E) & (F) Reconstructed image using the image in Fig. 8A with the algorithm described in section V(B). (G) Reconstructed image using the image in Fig. 8A with the algorithm described in section VI. Images B, E, and G are projected form the point marked with the white star. Image F is projected from the point marked with the black star.

We also show the results from reconstruction using the projection plane normal as reference (Fig. 8E). It is important to note that the distortion from reconstruction itself is more prominent with increase distance away from the point of projection as evident in Fig. 8F. However, because of the availability to project from any point on the surface, this enables a way to better resolve local features otherwise unavailable.

Last and the most general, we demonstrate also the reconstruction (Fig. 8F) without relying on a mathematical shape to describe the curved surface. Notice that the left boundary has better correction compared to Fig. 8B & E because this method makes no assumption on the shape. The size of the ommatidia is, however, of less consistency because of the piecewise reconstruction.

## VIII. DISCUSSION

Ellipsoidal geodesics problems have long been solved by Carl Jacobi [11]. His solution provides explicit formula for the computation of geodesics in elliptical coordinates. Although this solution minimizes the distance between 2 points on the surface of the ellipsoid, the contour of the geodesics is not necessary describable by a plane giving no easy way of planar projection, contrary to the demonstration here. Elliptical coordinates are also difficult to visualize thus the implementation is both error prone and non-intuitive. Employing the Jacobian solution will also limit the application to only elliptical structure, which is not the case here. The geometrical projection requires only 2 points, one being the point of projection, one being the center or any point along the projection plane normal. In a combinatorial manifold, if the user so choose to better describe the surface of interest, the center does not have to be defined as in section V(B).

The 2D mapping in section V relies on the fact that all 3D points are within close proximity to the mathematical surface. Therefore, a good fit is necessary to guarantee successful reconstruction. The quality of mapping is expected to be improved when employing a higher order description, but the choice of such depends on the particular situation and if small areas of misfit will significantly affect the final results. This is solved by not assuming any mathematical structure to describe the manifold as in section VI. However, as noted previously, the projection is not as smooth, because of the piecewise projection.

Most image analysis tools and platforms are designed to perform with 2D images. Although customized algorithm could be design to perform analysis 3D, it is both time consuming and computationally expensive. Reconstructing the 3D data to a 2D image without loss of information is ideal in optimizing efficiency and quality.

## ACKNOWLEDGMENT

This work was funded by the National Science Foundation (NSF) Grant 0965918 IGERT at UIUC: Training the Next Generation of Researchers in Cellular and Molecular Mechanics and Bio-Nanotechnology. The author would like to thank Dr. Minh Do and Dr. Taher Saif for technical comments.

## Footnotes

1 Department of Mechanical Science and Engineering, University of Illinois, Urbana, IL, 61801 USA

2 Department of Molecular Biosciences, Northwestern University, Evanston, IL, 60208 USA

